# Discovery of TMPRSS2 inhibitors from virtual screening

**DOI:** 10.1101/2020.12.28.424413

**Authors:** Xin Hu, Jonathan H. Shrimp, Hui Guo, Miao Xu, Catherine Z. Chen, Wei Zhu, Alexey Zakharov, Sankalp Jain, Paul Shinn, Anton Simeonov, Matthew D. Hall, Min Shen

## Abstract

The SARS-CoV-2 pandemic has prompted researchers to pivot their efforts to finding antiviral compounds and vaccines. In this study, we focused on the human host cell transmembrane protease serine 2 (TMPRSS2), which plays an important role in the viral life cycle by cleaving the spike protein to initiate membrane fusion. TMPRSS2 is an attractive target and has received attention for the development of drugs against SARS and MERS. Starting with comparative structural modeling and binding model analysis, we developed an efficient pharmacophore-based approach and applied a large-scale *in silico* database screening for small molecule inhibitors against TMPRSS2. The hits were evaluated in the TMPRSS2 biochemical assay and the SARS-CoV-2 pseudotyped particle (PP) entry assay. A number of novel inhibitors were identified, providing starting points for further development of drug candidates for the treatment of COVID-19.

---

Severe acute respiratory syndrome coronavirus 2 (SARS-CoV-2) is the virus responsible for the ongoing coronavirus disease 2019 (COVID-19) pandemic.^1^ It is a novel betacoronavirus from the same family as SARS-CoV and Middle East respiratory syndrome (MERS).^2^ SARS-CoV-2 produces clinical symptoms that include fever, dry cough, sore throat, dyspnea, headache, pneumonia with potentially progressive respiratory failure owing to alveolar damage, and even death.^3^ Recently it has also been reported that COVID-19 is associated with the amplified incidence of thrombotic events, which contribute to severe coagulation in different organs such as lung, brain, liver, heart, kidney and further lead to multi-organ failure and death.^4, 5^ SARS-CoV-2 is a global pandemic with millions of documented infections worldwide, and over 1 million deaths reported by World Health Organization (WHO). Since no antiviral drug or vaccine existed to treat or prevent SARS-CoV-2, potential therapeutic strategies that are currently being evaluated predominantly stem from previous experience with treating SARS-CoV, MERS-CoV, and other emerging viruses.^6-8^ Most nations are primarily making efforts to prevent the further spreading of this potentially deadly virus by implementing preventive and control strategies.

A number of processes are considered essential to the viral lifecycle and therefore provide a significant number of targets for inhibiting viral replication. The screening of anti-COVID-19 drugs by using the clinical and approved compounds can greatly shorten the research and development cycle. Some screening campaigns of approved drug libraries and pharmacologically active molecules have been conducted for the discovery of SARS-CoV-2 inhibitors. Antiparasitic drugs like chloroquine and its derivative hydroxychloroquine have shown antiviral activity against SARS-CoV-2 in an *in vitro* cytopathic assay and had an Emergency Use Authorization awarded and rescinded by the US FDA.^9, 10^ Zn^(2+)^ was found to inhibit coronavirus through blocking the initiation step of equine arteritis virus RNA synthesis *in vitro* and zinc ionophores inhibit the replication of coronavirus in cell culture.^11^ Besides having direct antiviral effects, chloroquine and hydroxychloroquine specifically target extracellular zinc to intracellular lysosomes where it interferes with RNA-dependent RNA polymerase activity and inhibits coronavirus replication.^12^ The broad-spectrum antibacterial macrolide azithromycin, an inhibitor of bacterial protein synthesis, also has significant antiviral properties, and it decreased the coronavirus infection in cell culture.^13^ The antiviral lopinavir, a protease inhibitor, has also shown inhibitory activity of SARS-CoV-2 infection in Vero E6 cells.^14^ The experimental drug remdesivir, a nucleoside analog, originally developed against other viruses, has been approved by the FDA as treatment for COVID-19.^9, 15^ Despite numerous biochemical and cell-based drug repurposing screening of drug compound libraries that have identified a number of potent antivirals targeting various stages of the viral life cycle, the development of effective intervention strategies relies on the knowledge of molecular and cellular mechanisms of coronavirus infections, which highlights the significance of studying virus–host interactions at the molecular level to identify targets for antiviral intervention and to elucidate critical viral and host determinants that are decisive for the development of severe disease.^16^

Cell entry of coronaviruses depends on binding of the viral spike (S) proteins to human cellular receptors and on S protein priming by host cell proteases. It has been demonstrated that SARS-CoV-2 uses the human host cell angiotensin-converting enzyme 2 (ACE2) as the entry receptor and the transmembrane protease serine 2 (TMPRSS2) for S protein priming.^17, 18^ TMPRSS2 has an extracellular protease domain capable of cleaving a spike protein domain to initiate membrane fusion. Considering the vital role played by TMPRSS2 in the viral life cycle, this protease has received great attention to be used as a potential target to inhibit viral entry into host cells.^19-21^ Bestle *et al* used an antisense peptide-conjugate to interfere with splicing resulting in the expression of a truncated, enzymatically dead TMPRSS2. This is the most targeted experiment showing that KD of enzymatic activity lowered cytopathic effect, viral spread and replication of SARS-CoV-2 in Calu-3 cells.^22^ More recently, Hoffmann *et al* showed that camostat mesylate and its metabolite can lower viral entry in Calu-3 cells where there is a TMPRSS2-dependent viral entry. Additionally, they show from single-cell RNA-Seq datasets that 53% of ACE2 expressing cells co-express TMPRSS2 in lungs, indicating that the tissue-specific TMPRSS2 is the dominant activating protease in lung.^23^

Previous basic and clinical research on coronaviruses has led to the identification of many potential drug targets and determination of their X-ray crystal structures. Structure-based drug design by virtual screening and molecular docking studies has become a valuable primary step in the identification of novel lead molecules for the potential treatment of COVID-19.^24^ In this study, we performed a comprehensive structural modeling and binding site analysis of the serine protease TMPRSS2, followed by a structure-based virtual screening against NCATS library consisting of up to 200,000 drug-like compounds designed from diverse chemical space. After an extensive post-docking analysis in combination with clustering analysis and visual inspection, 350 compounds were selected for experimental validation based on their binding free energy, consensus docking score, and binding interactions with key residues surrounding the active site. The selected hits were evaluated in the TMPRSS2 biochemical assay and the SARS-CoV-2-S pseudotyped particle (PP) entry assay.^25, 26^ A number of novel inhibitors were identified, providing a starting point for further development of therapeutic drug candidates for COVID-19.

## Materials and methods

### Homology modeling of TMPRSS2

Amino acid sequence of human TMPRSS2 was obtained from the UniProtKB (access number O15393). The protein consisting of 492 amino acid residues is a zymogen cleaved at Arg-255. The protease domain (255-492) is located at the C terminus of the extracellular region, which also contains a SRCR activation domain (150-242) and a short LDLRA domain (113-148). We modeled the 3D structure of TMPRSS2 SRCR and peptidase S1 domain (150-492) using the threading program I-TASSER.^27^ The transmembrane trypsin-like serine protease hepsin (also known as TMPRSS1, gene *HPN*) which shares 35% of sequence identity with TMPRSS2, was used as the template structure (PDB code 1Z8G).^28^ An overlay of TMPRSS2 structure model with template hepsin and other two related serine proteases coagulation factor Xa (PDB 1KSN)^29^ and urokinase (PDB 1F92)^30^ is shown in **Figure S1**. The generated structure was validated using the tools (PROCHECK, ERRAT, Verify3D, PROVE and WHATCHECK) in the SAVES server (https://servicesn.mbi.ucla.edu/SAVES/). The modeled structure passed in all programs, indicating a good quality overall. Finally, the 3D structure of TMPRSS2 was refined by thorough energy minimization and MD simulations as described below.

### Molecular docking and MD simulations

Three TMPRSS2 inhibitors nafamostat, camostat, and gabexate were used in this study. Nafamostat mesylate is an approved anticoagulant in Asian countries and currently in Phase 3 clinical trials for COVID-19.^31^ Camostat mesylate was approved for human use in Japan for the treatment of chronic pancreatitis and postoperative reflux esophagitis. It is a broad spectrum serine protease inhibitor targeting coronavirus and filovirus entry.^32^ Gabexate is an investigational drug and serine protease inhibitor used to prevent blood clots and reduce the production of inflammatory cytokines.^33^ The activities of these inhibitors against TMPRSS2 have been confirmed in our recent report, with IC_50_ of 0.27 nM, 6.2 nM, and 130 nM, respectively.^25^

Consensus docking studies of these small molecule inhibitors to the modeled structure of TMPRSS2 were performed using MOE Dock in the Molecular Operating Environment program^34^ and AutoDock Vina.^35^ Prior to docking, the 3D structure of TMPRSS2 was prepared using the Structure Preparation module in MOE. The ligand induced fit docking protocol in MOE Dock was applied and binding affinity was evaluated using the GBVI/WSA score. The default parameters in AutoDock Vina were used with a grid box centered on the O atom of the side chain of catalytic residue Ser441. The size of grid box was defined by 20 × 20 × 20 Å to encompass the entire active site of the protein. The top-ranked 10 poses from MOE Dock and AutoDock Vina were retained and visually inspected. The consensus binding models with lowest binding energies were selected for further MD simulations.

MD simulations of the TMPRSS2 in the apo form and inhibitor-bound complexes were conducted using the AMBER18 package.^36^ The protein was protonated at pH 7. The catalytic residue His296 was protonated as N-δ form so that it formed H-bonding with Ser441 and Asp345. The solvated protein systems were subjected to a thorough energy minimization prior to MD simulations by first minimizing the water molecules while holding the solute frozen (1000 steps using the steepest descent algorithm), followed by 5,000 steps of conjugate gradient minimization of the whole system to relax the system. Periodic boundary conditions were applied to simulate a continuous system. The particle mesh Ewald (PME) method was employed to calculate the long-range electrostatic interactions. The simulated system in explicit solvate was first subjected to a gradual temperature increase from 0 K to 300 K over 100 ps, and then equilibrated for 500 ps at 300 K, followed by a production run of 100 ns. Trajectory analysis and MM-GBSA binding free energy calculations were performed using the cpptraj and MMPB/GBSA module in the AmberTools18.^36^

### Pharmacophore-based virtual screening

A stepwise virtual screening (VS) protocol combining ligand- and structure-based approach was employed to screen the NCATS’s in-house screening libraries consisting of nearly 200,000 drug-like compounds.^37^ Pharmacophore models were generated based on the predicted binding interactions of TMPRSS2 with inhibitor camostat, nafamostat and gabexate using MOE. Four pharmacophoric features were included: 1) an Don2 projected H-bond donor feature placed on the sidechain of Asp435 in the S1 pocket; 2) an Acc2 projected H-bond acceptor feature placed on the N atom of sidechain of Gln438; 3) an hydrophobic centroids Hyd feature matching hydrophobic interactions at the S1’ hydrophobic region mainly formed by Val275, Val280, and Leu302; 4) an Don2 projected H-bond donor feature placed on the sidechain of Glu299. The first two pharmacophores were selected as essential features and database searching was conducted with partial match mode using MOE. To enrich the structural diversity of potential hits, the 3D shape-based searching was applied using ROCS.^38^ The predicted binding conformations of three inhibitors were used as queries.

A total of 20,000 hits were extracted from the pharmacophore and ligand-based database searching, followed by docking to the active site of TMPRSS2 using MOE dock and AutoDock Vina with default parameters.^35^ The top-ranked 2000 compounds from each docking were retained for consensus scoring and binding mode analysis. Structural clustering was performed using MOE. All compounds from each cluster and singletons were visually inspected. Finally, 350 compounds were selected based on: 1) structural representative of each cluster, 2) predicted binding energy, 3) H-bond interaction with key residues at each binding site, 4) promiscuous compounds with potential undesirable functionalities and PAINS alert were generally discarded.

### TMPRSS2 biochemical assay

A TMPRSS2 biochemical assay has been developed to evaluate the activity of compounds against TMPRSS2.^25^ The assay was performed according to the assay protocol: to a 1536-well black plate was added Boc-Gln-Ala-Arg-AMC substrate (20 nL) and test compound (20 nL in DMSO) using an ECHO 655 acoustic dispenser (LabCyte). To that was dispensed TMPRSS2 diluted in assay buffer (50 mM Tris pH 8, 150 mM NaCl, 0.01% Tween20) using a BioRAPTR (Beckman Coulter) to give a total assay volume of 5 µL. Following 1 h incubation at room temperature, fluorescence was measured using the PHERAstar with excitation at 340 nm and emission at 440 nm.

### Fluorescence counter assay

This counter-assay was performed as described previously.^25^ To a 1536-well black plate (Corning Cat #3724) was added 7-amino-4-methylcoumarin (20 nL) and inhibitor or DMSO (20 nL) using an ECHO 655 acoustic dispenser (LabCyte). To that was added assay buffer (50 mM Tris pH 8, 150 mM NaCl, 0.01% Tween20) to give a total reaction volume of 5 µL. Detection was done using the PHERAstar with excitation: 340 nm and emission: 440 nm. Fluorescence was normalized relative to a negative control 7-amino-4-methylcoumarine. An inhibitor causing fluorescence quenching would be identified as having a concentration-dependent decrease on AMC fluorescence.

### SARS-CoV-2-S pseudotyped particle entry assays

SARS-CoV-2 Spike pseudotyped particle (PP) entry assay was performed as previously described.^39^ In short, 40,000 cells/well Calu3 cells (Cat #HTB-55, ATCC) were seeded in white, solid bottom 96-well microplates (Greiner BioOne) in 100 µL/well media, and incubated at 37 °C with 5% CO2 overnight (∼16 h). Then, the supernatant was removed, and compounds were added as 50 μ l/well, 2x solutions in media. Cells were incubated with compounds for 1 h at 37 °C with 5% CO2, before 50 µL/well of SARS-CoV-2-S pseudotyped particles (PP) was added. The PP pseudotyped SARS-CoV-2 spike from the South African variant B.1.351 (Cat #CB-97100-154, Codex Biosolutions) with a C-terminal 19 amino acid deletion using the murine leukemia virus system as previously described.^40^ The plates were then spinoculated by centrifugation at 1500 rpm (453 xg) for 45 min, and incubated for 48 h at 37 °C 5% CO2 to allow cell entry of PP and expression of luciferase reporter. After the incubation, the supernatant was removed and 100 µL/well of Bright-Glo Luciferase detection reagent (Promega) was added to assay plates and incubated for 5 min at room temperature. The luminescence signal was measured using a PHERAStar plate reader (BMG Labtech). Data was normalized with wells containing SARS-CoV-2-S PPs as 100%, and wells containing control bald PP as 0%. A cytotoxicity counter screen was performed using the same protocol without the addition of PP, with an ATP content assay kit (CellTiterGlo, Promega).

## Results

### Binding interaction of TMPRSS2 inhibitor

TMPRSS2 shares 35% of sequence identity with the transmembrane trypsin-like serine protease hepsin. The modeled structure of TMPRSS2 generated from I-TASSER had a C-score of -0.10 and TM-score of 0.70 which indicated a good quality modeled structure.^27^ Similar to the template hepsin and other trypsin-like proteases, TMPRSS2 shares a common structural fold with a conserved triad residues Ser441, His296, and Asp345 at the active site for catalytic activity.^41, 42^ A predicted oxyanion hole is formed by conserved residues Gly439 and Gln438 at the active site for the catalytic process and a highly hydrophilic S1 pocket with a conserved Asp435 which is essential for substrate and inhibitor binding.

The predicted binding models of camostat, nafamostat, and gabexate to the active site of TMPRSS2 are shown in **Figure 1**. The three small molecule inhibitors adopted a similar binding mode at the active site of TMPRSS2. While the ester group of the guanidinobenzonate moiety of camostat and nafamostat binds at the triad catalytic site interacting with Ser441 and Gln438, the guanidinium head points into S1 pocket forming H-bonding and ion interactions with Asp435. The same binding models have been reported by Hempel *et al*, showing that the ester group resembled the substrate peptide bond by forming a covalent bond to the catalytic Ser441, and the high potency of nafamostat can be explained from a greater stability of its Michaelis complex.^43^ Gabexate binds in a similar manner with the essential guanidinium head interacting with Asp435 in the S1 pocket and the ester group forming hydrogen bonds with Ser441 and Gln438, whereas the benzoic ester group points to the hydrophobic region at S1’ site, same as observed with camostat binding. The naphthylamidinium of nafamostat forms an H-bonding interaction with Glu299. Interestingly, a reverse binding mode of nafamostat was found that the amidinium head bound into the S1 pocket interacting with Asp435, which showed comparable binding free energy. Such an inverted binding orientation of nafamostat has been investigated by Hempel *et al*, suggesting that it may also be reactive with TMPRSS2 at the catalytic site.^43^

**Figure 1.**
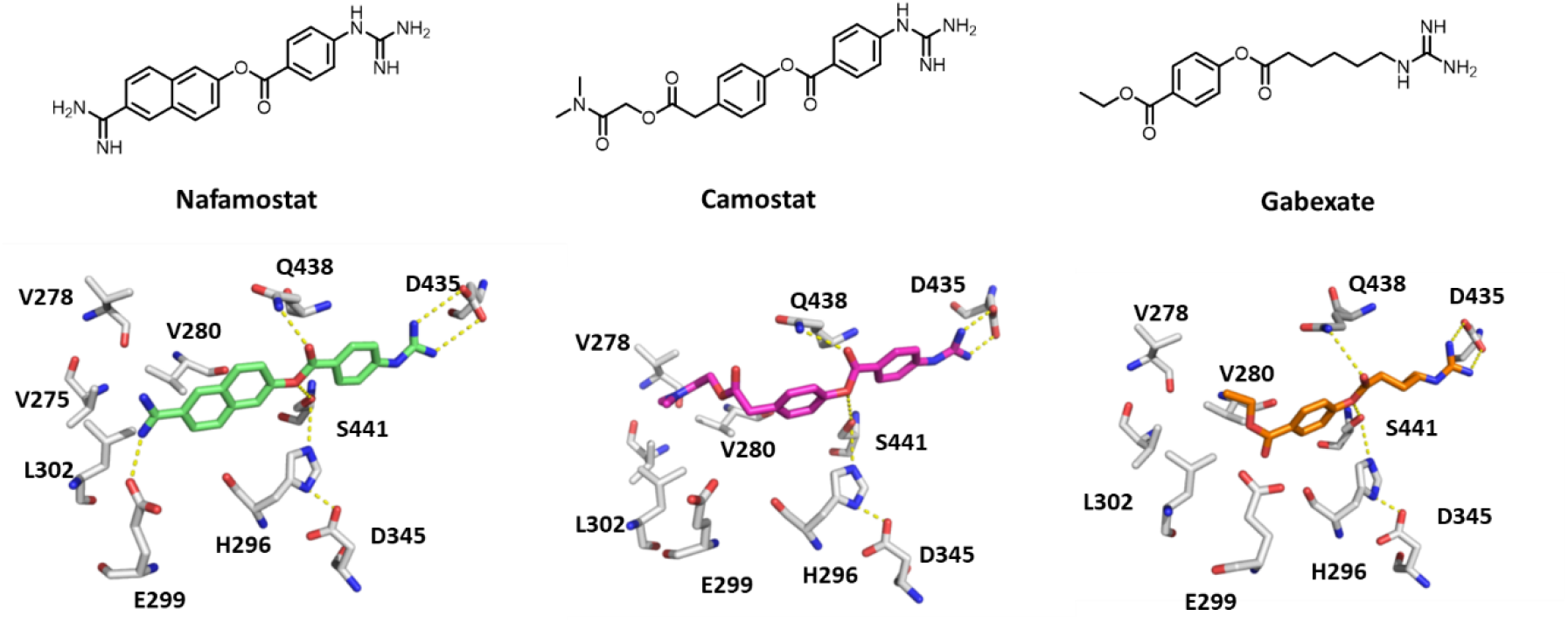
Known TMPRSS2 inhibitors and predicted binding models at the active site of TMPRSS2.

To gain insight into the binding interactions of these inhibitors with TMPRSS2, we performed MD simulations of the inhibitor-bound complexes as compared to the protein in the apo state (**Figure 2A**). The loops surrounding the active site exhibited greater dynamics in the apo form, but were generally stabilized in the inhibitor binding complex over the time course of 100-ns simulations. The hydrogen bonds between the guanidinium head and residues Asp435 in the S1 pocket remained stable in the binding complexes (**Figure S2** and **Figure S3**). Gabexate binding complex was more dynamical in the MD simulations, which may explain its weaker activity as compared to camostat and nafamostat. Notably, residues Glu299 and Gln438 were found to be rather flexible in the MD simulations. These residues are in the active site adjacent to the catalytic triad, which likely play an important role in facilitating the substrate recognition as well as inhibitor binding.

**Figure 2.**
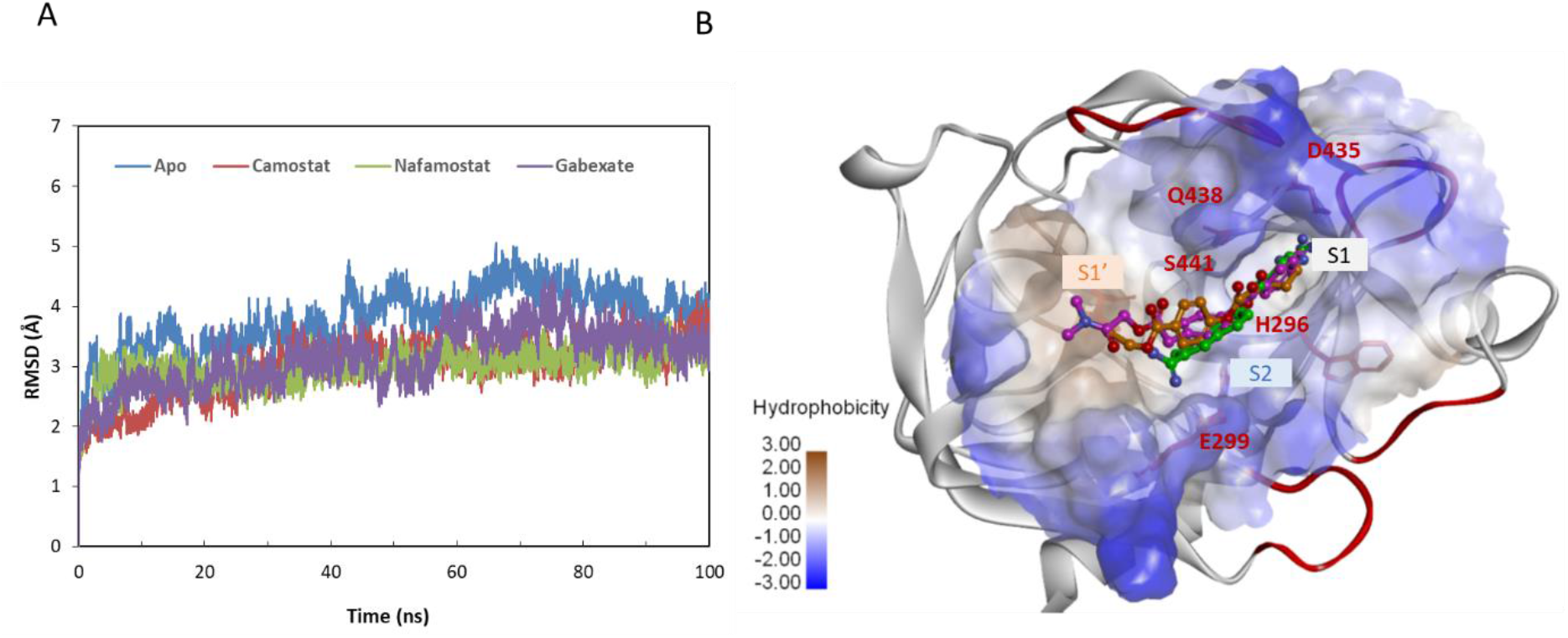
(A) MD simulations of TMPRSS2 in the apo and inhibitor-bound complexes. (B) Inhibitors camostat (magenta), nafamostat (green) and gabexate (dark yellow) bound at the active site of TMPRSS2. The protein surface is rendered in color of hydrophobicity. Dynamics loops surrounding the active site are shown in red. Key residues and the hydrophobic pocket used in pharmacophore model are labeled.

### Pharmacophore-based virtual screening

Based on the binding model analysis we developed a pharmacophore model and applied it in virtual screening for novel inhibitors of TMPRSS2. Four pharmacophores were derived from the predicted binding interactions of the three known inhibitors (**Figure S4**). Two binding elements were selected as essential features: first is an H-bonding interaction between Asp435 and a functional head in the S1 pocket; second is a core H-bonding interaction with Gln438, which facilitates inhibitor binding with the triad residues at the catalytic site. The other two pharmacophoric features included the hydrophobic interactions at the S1’ pocket mainly formed by Val275, Val280, and Leu302 which serves as an extended region to accommodate variable groups for enhanced binding affinity, the other one is a binding interaction with Glu299 which provides a key to position the inhibitor into the binding pocket.

Virtual screening was performed in a stepwise way combined with pharmacophore-based searching, 3D-shape-based mapping, and structure-based docking (**Figure 3**). An in-house collection of nearly 200,000 drug-like compounds were virtually screened. The pharmacophore model was firstly applied and compounds matching at least two pharmacophores were extracted. To enrich the structural diversity of potential hits, the 3D shape-based searching was applied using the predicted binding conformations of three inhibitors as a query. A total of 20,000 compounds were assembled from the pharmacophore- and ligand-based search, followed by structure-based docking to the active site of TMPRSS2 for binding interaction evaluation. To prioritize the hits generated from docking, several post-processing approaches were applied including docking pose analysis, structural clustering, re-scoring, and promiscuity filtering. Ultimately, ∼350 compounds were selected based on the predicted binding energy, key interactions within the catalytic triad and functional head in the S1 pocket, the novelty of scaffold and chemotypes. Promiscuous compounds (i.e., those shown to have high hit rates across HTS assays experimentally screened at NCATS) with potential undesirable functionalities and PAINS alerts were generally discarded.

**Figure 3.**
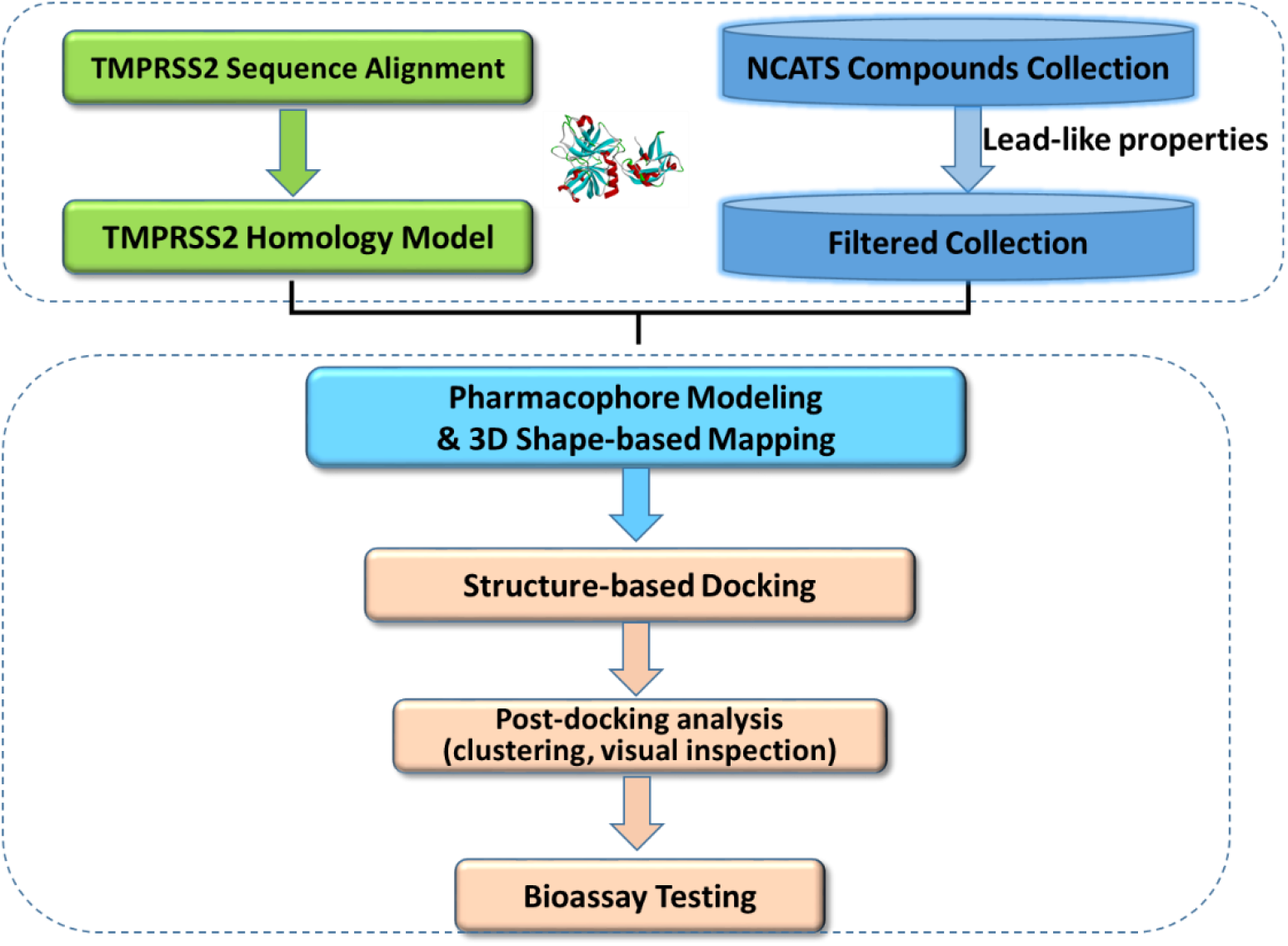
Flowchart of virtual screening.

### TMPRSS2 inhibitors identified from VS

All selected compounds were evaluated in the TMPRSS2 enzymatic assay, a fluorogenic biochemical assay we reported recently.^25^ A counter assay was conducted to detect all positive compounds with fluorescence quenching properties that suppress the fluorescence signal generated by the protease activity on the fluorogenic substrate. Of 350 compounds tested, 27 hits showed activity against TMPRSS2 with efficacy greater than 30% and an IC_50_ ranging from 1 µM to 30 µM. The active compounds were further re-tested in the biochemical and counter assay at 11 concentrations in triplicate (**Table 1 and Figure S5). Figure 4** showed the top hits with efficacy greater than 50% and IC_50_ lower than 10 µM. The three inhibitors camostat, nafamostat and gabexate were used as positive controls and all showed the same inhibitory activities as previously reported.^25^ The best VS hits of NCGC00378763 (otamixaban) and NCGC00421880 showed IC_50_ of 0.62 µM and 0.88 µM. A selective inhibitor of urokinase, UKI-1 (NCGC00522442), showed IC_50_ of 2.20 µM with 100% inhibition. While most of these identified inhibitors possess a benzoamidinium head group, they are structurally diverse with distinct scaffolds and molecular properties different from nafamostat and camostat.

**Table 1.**
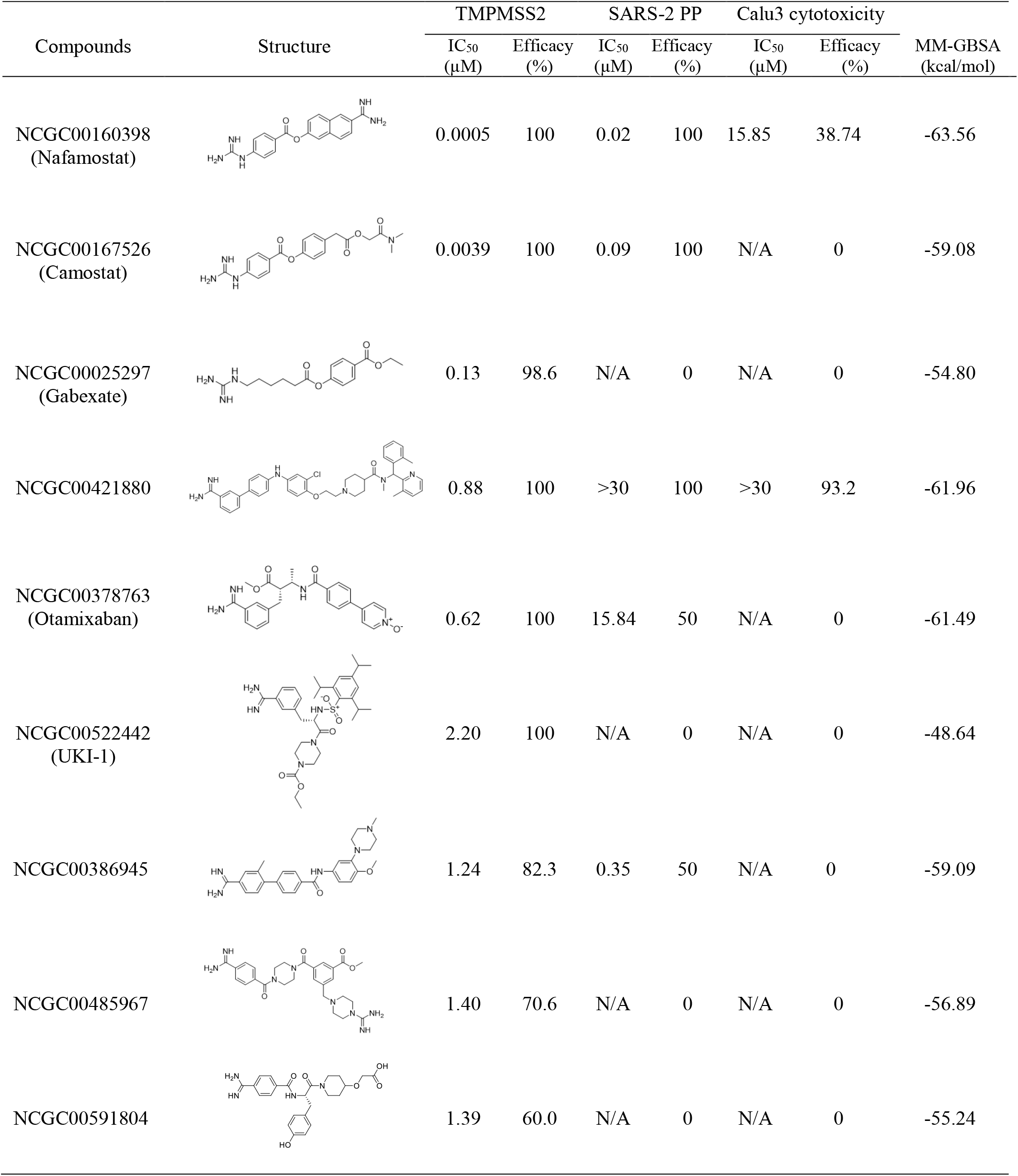
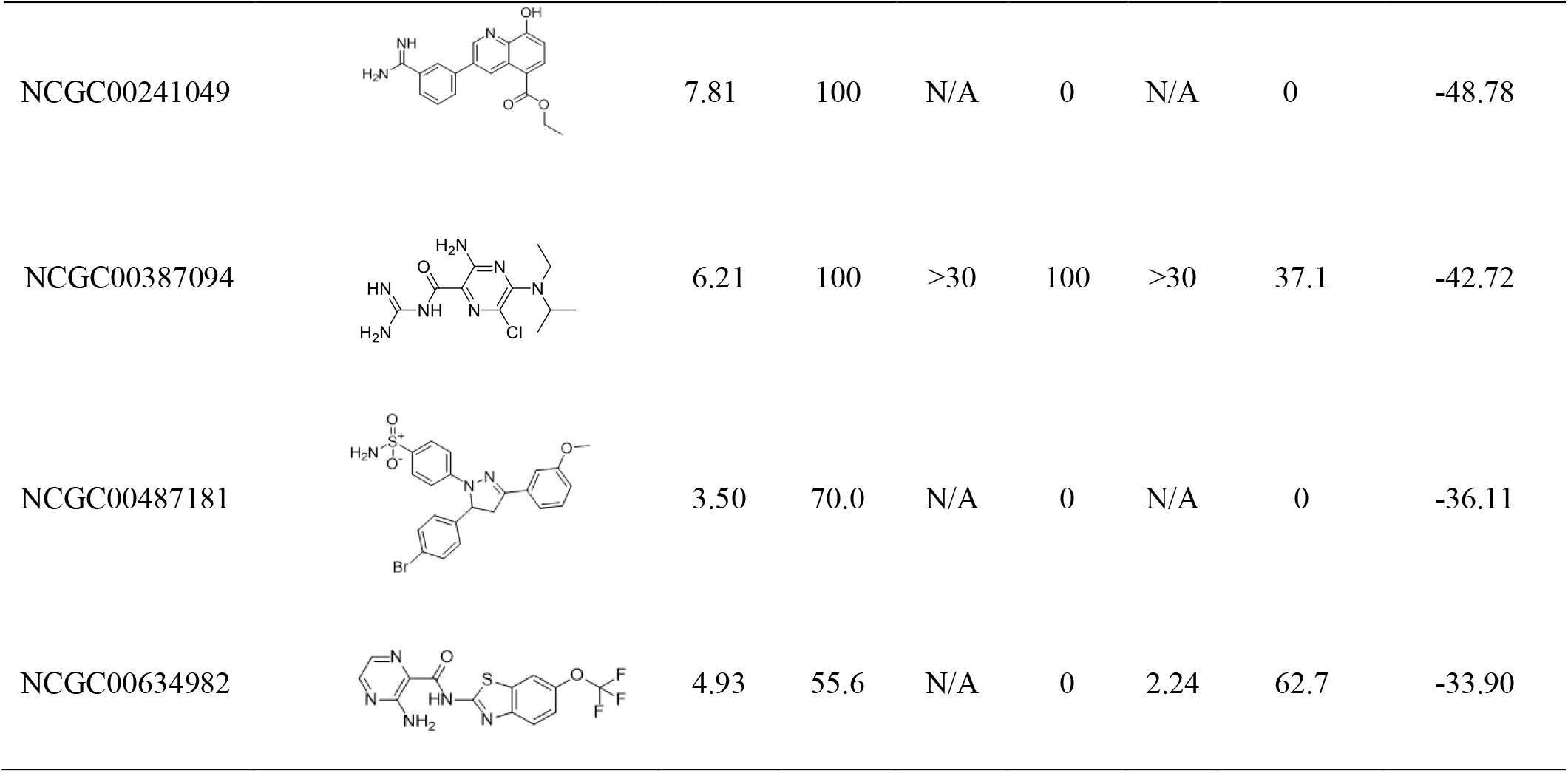
Activity of TMPRSS2 inhibitors in the enzyme assay and the SARS-COV-2 PP entry assay. The MM-GBSA binding free energy was estimated based on the predicted binding model from docking.

| Compounds | Structure | TMPRSS2 |  | SARS-2 PP |  | Calu3 cytotoxicity |  | MM-GBSA (kcal/mol) |
| --- | --- | --- | --- | --- | --- | --- | --- | --- |
|  |  | IC<sub>50</sub> (μM) | Efficacy (%) | IC<sub>50</sub> (μM) | Efficacy (%) | IC<sub>50</sub> (μM) | Efficacy (%) |  |
| NCGC00160398<br>(Nafamostat) |  | 0.0005 | 100 | 0.02 | 100 | 15.85 | 38.74 | -63.56 |
| NCGC00167526<br>(Camostat) |  | 0.0039 | 100 | 0.09 | 100 | N/A | 0 | -59.08 |
| NCGC00025297<br>(Gabexate) |  | 0.13 | 98.6 | N/A | 0 | N/A | 0 | -54.80 |
| NCGC00421880 |  | 0.88 | 100 | >30 | 100 | >30 | 93.2 | -61.96 |
| NCGC00378763<br>(Otamixaban) |  | 0.62 | 100 | 15.84 | 50 | N/A | 0 | -61.49 |
| NCGC00522442<br>(UKI-1) |  | 2.20 | 100 | N/A | 0 | N/A | 0 | -48.64 |
| NCGC00386945 |  | 1.24 | 82.3 | 0.35 | 50 | N/A | 0 | -59.09 |
| NCGC00485967 |  | 1.40 | 70.6 | N/A | 0 | N/A | 0 | -56.89 |
| NCGC00591804 |  | 1.39 | 60.0 | N/A | 0 | N/A | 0 | -55.24 |
| NCGC00241049 |  | 7.81 | 100 | N/A | 0 | N/A | 0 | -48.78 |
| NCGC00387094 |  | 6.21 | 100 | >30 | 100 | >30 | 37.1 | -42.72 |
| NCGC00487181 |  | 3.50 | 70.0 | N/A | 0 | N/A | 0 | -36.11 |
| NCGC00634982 |  | 4.93 | 55.6 | N/A | 0 | 2.24 | 62.7 | -33.90 |

**Figure 4.**
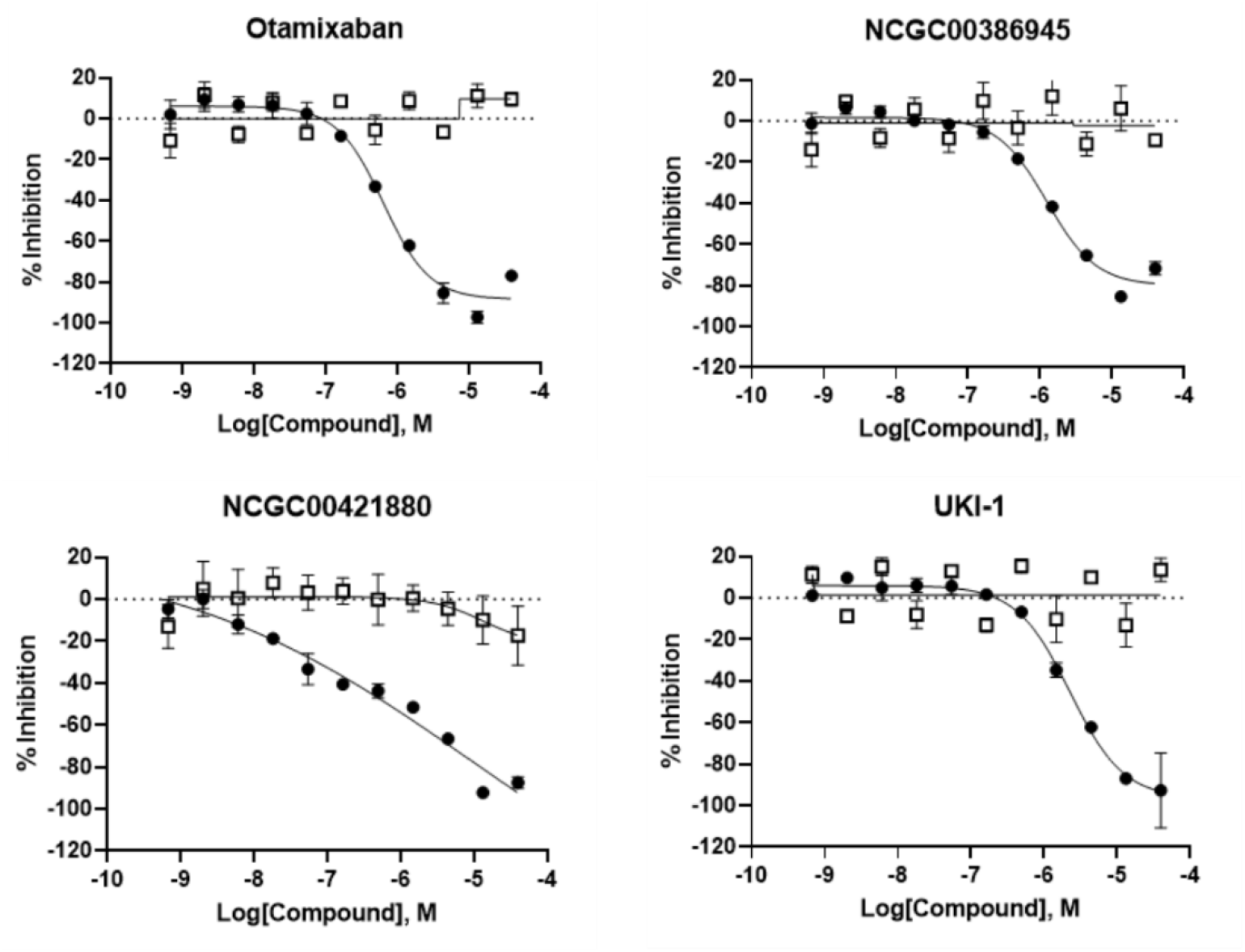
Activities of identified inhibitors in the TMPRSS2 enzyme assay (solid circle) and fluorescence counter screen (empty square).

We further tested the inhibitors in a SARS-CoV-2 spike pseudotyped particle (PP) entry assay in Calu-3 cells to evaluate their inhibitory activities on viral entry. While SARS-CoV-2 spike could utilize multiple cellular proteases for entry, entry into Calu-3 cell has been shown to be predominantly through the TMPRSS2-mediated pathway.^21^ The PPs pseudotype the South African variant (B.1.351) spike protein bearing the mutations L18F, D80A, D215G, del242-244, K417N, E484K, N501Y, D614G, A701V. These mutations do not affect the S2’ cleavage site recognized by TMPRSS2. A cytotoxicity counter assay was conducted using the same protocol without the addition of PP. Nafamostat and camostat showed high potency and efficacy in the SARS-CoV-2 PP assay, however, gabexate did not exhibit activity on viral entry inhibition (**Figure 5**). Of the 10 VS inhibitors tested, only otamixaban and NCGC00386945 displayed activity with about 50% efficacy in the PP entry assay, whereas NCGC00421880 and NCGC00387094 showed SARS-CoV-2 inhibition as well as cytotoxicity in the counter assay (**Table 1**). No inhibition activity was observed with other inhibitors in the cell-based assay most likely because of their weak activities against TMPRSS2. We also evaluated three TMPRSS2 inhibitors of avoralstat, PCI-27483 and antipain which have been reported recently.^44^ While avoralstat displayed comparable inhibition activity to camostat, other two protease inhibitors showed about 60% efficacy activity in our PP entry assay (**Figure S6)**.

**Figure 5.**
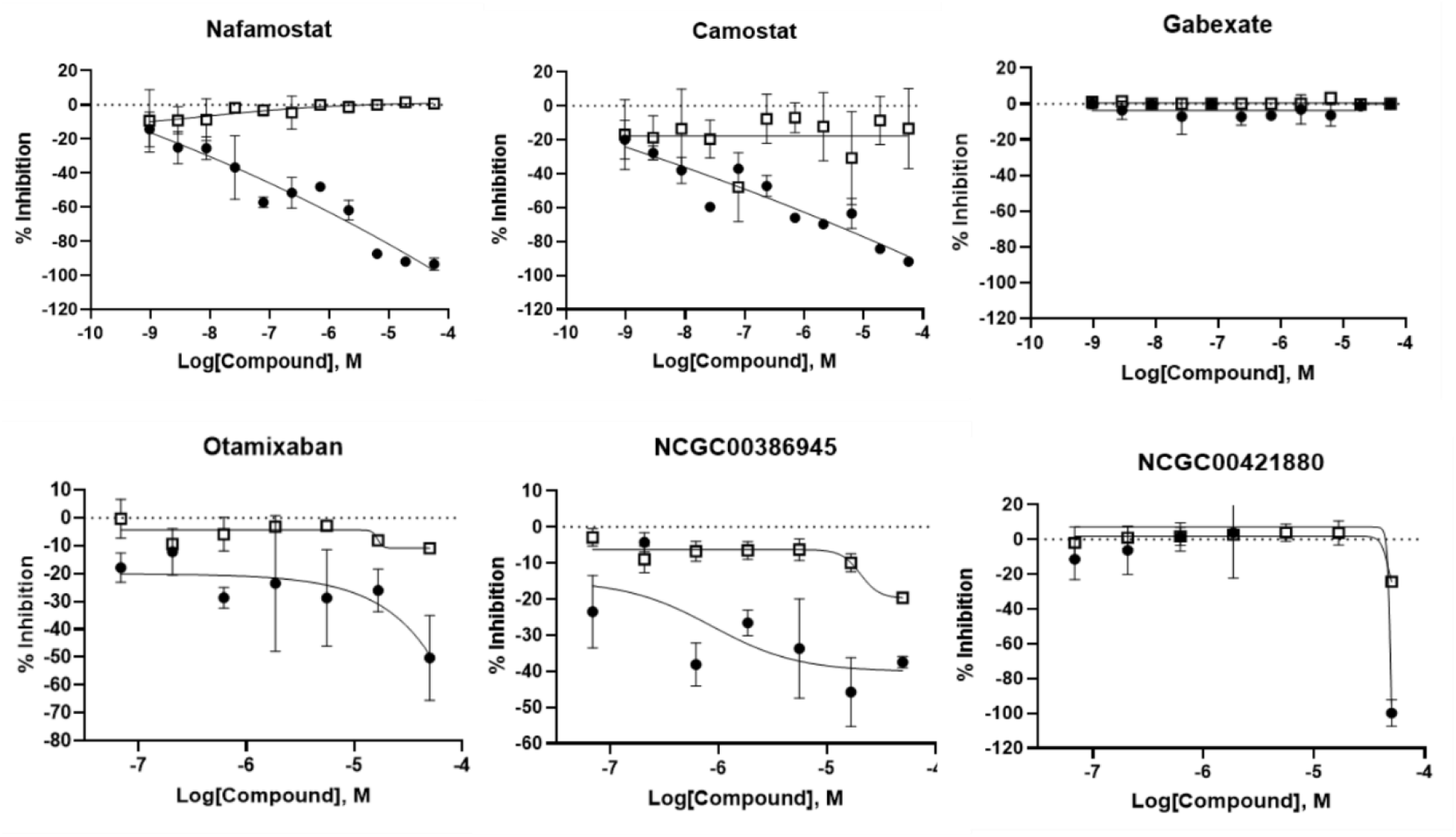
Activities of TMPRSS2 inhibitors in the SARS-COV-2-S PP entry assay (solid circle) and cytotoxicity counter screen (empty square).

### Binding interaction of identified inhibitors at the active site of TMPRSS2

Finally, we performed docking studies and MD simulations to probe the binding interaction of these identified inhibitors with TMPRSS2. **Figure 6** showed the predicted binding model of the top four inhibitors bound at the active site of TMPRSS2. As expected, these small molecules utilized the benzoamidinium head group to engage key H-bonding interactions with Asp435 in the S1 pocket, while the tail group was mainly positioned at the S1’ hydrophobic pocket. Other inhibitors showed similar binding interactions to camostat and nafamostat at the catalytic site and S1’ pocket (**Figure S7)**. MD simulations showed that these inhibitors remained stable at the active site of TMPRSS2. However, unlike camostat and nafamostat binding which form stable H-bonding interactions between the guanidinobenzoyl moiety and the catalytic triad residues for covalent acyl-enzyme complex formation, these non-covalent inhibitors do not have such a functionality that makes favorable H-bond interactions at the catalytic site. These inhibitors bound at the active site predominately through hydrophobic interaction at S1’ and S2 pocket. The calculated binding free energies based on the predicted binding model of these non-covalent inhibitors were generally consistent with the experimental data (**Table 1**). Notably, docking analysis showed that otamixaban adopted another distinct binding mode with the pyridine-oxide group pointing to the S2 and S4 binding pocket. The calculated binding free energies of the two binding conformations with TMPRSS2 were -61.49 and -67.69 kcal/mol, suggesting that the second binding mode is likely more favorable (**Figure S7)**. Such a binding mode of otamixaban was also observed in the structure with coagulation factor Xa.^29^ In addition, the bulky triisopropylphenylsulfonyl group of UKI-1 was also occupied at the S2 pocket by mainly interacting with Trp461. TMPRSS2 possesses several polar residues including a unique Lys342 at this site, which makes it possible to further optimize the binding interactions to improve the potency and selectivity to TMPRSS2.

**Figure 6.**
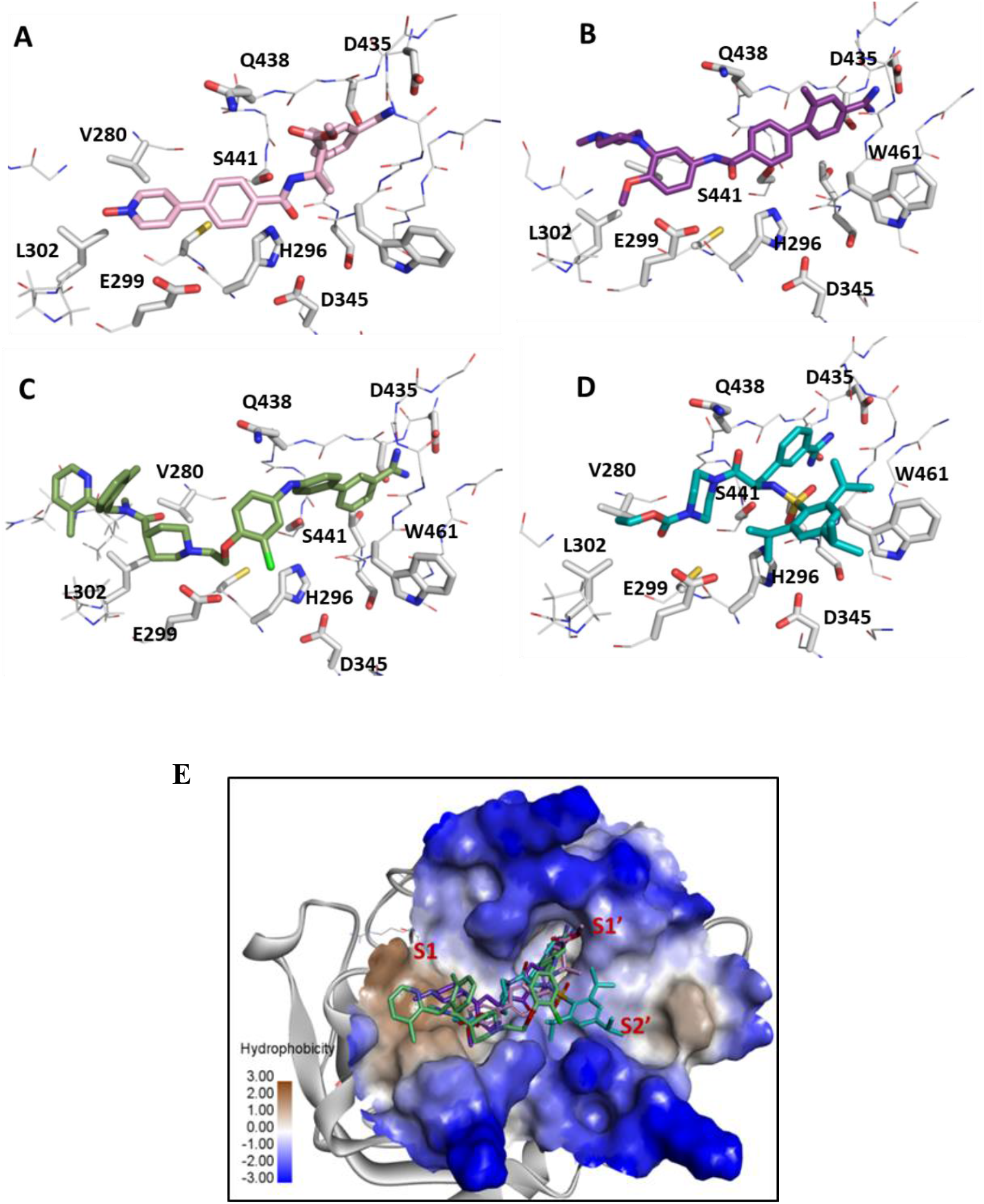
Predicted binding model of inhibitors (A) otamixaban, (B) NCGC00386945, (C) NCGC00421880, and (D) UKI-1. (E) An overlay of four inhibitors bound at the active site of TMPRSS2 is shown. Protein surface is rendered in hydrophobic representation.

## DISCUSSION

The host serine protease TMPRSS2 is an attractive target for drug development against SARS-CoV-2. However, unlike viral proteases such as 3Clpro which has been extensively investigated recently and a large number of inhibitors have been reported,^45^ TMPRSS2 has been less studied and very few compounds have been evaluated in drug re-purposing screening. The enzyme is membrane associated, making it more challenging to express recombinant protein and study *in vitro*. Although camostat and nafamostat have progressed into clinical trials for the treatment of COVID-19, these inhibitors also show potent activity against many other trypsin-like serine proteases such as the plasma trypsin-like proteases, plasmin and FXIa.^25^ Therefore, there is an unmet need for novel and selective candidates of TMPRSS2 inhibition for further drug development. Additionally, in theory the TMPRSS2 inhibitors have potential applications for emerging variants of SARS-CoV-2 along with potential future viral outbreaks beyond SARS-CoV-2.

We began the virtual screening campaign by iteratively searching and docking nearly 200,000 compounds to the active site of TMPRSS2, followed by testing 350 compounds in the enzyme assay. The active compounds were further tested in cell-based PP assay to evaluate their activities on spike-mediated entry inhibition. Several interesting inhibitors were identified from the structure-based VS. Otamixaban is an experimental anticoagulant direct factor Xa inhibitor that was investigated for the treatment of acute coronary syndrome but its development was terminated in a Phase III clinical trial due to poor performance.^46^ Otamixaban exhibited potent activity with an IC_50_ of 0.62 µM against TMPRSS2 and efficient activity in the viral entry assay, indicating that it is a promising candidate to further optimization and drug re-purposing. NCGC00386945 is a novel inhibitor that was originally developed as a selective 5-HT1D antagonist.^47^ It showed activity against TMPRSS2 with an IC_50_ of 1.24 µM, as well as 50% efficacious activity on the viral entry inhibition. Similar to otamixaban, this novel chemotype of TMPRSS2 inhibitor with drug-like properties is structurally appealing for further lead optimization (**Table S1**). As shown in the predicted binding model, substitution of the methyl group at the benzoamidinium head with a polar group may engage an H-bonding interaction with Gln438, thus significantly improve the binding affinity and selectivity to TMPRSS2. UKI-1 is a selective inhibitor of trypsin and uPA.^48^ While it displayed a value of IC_50_ within single-digit micromolar against TMPRSS2, it did not show activity in our PP entry assay. This was also observed with inhibitor Gabexate, which had an IC_50_ below micromolar in the TMPRSS enzyme assay but was inactive in the viral entry cell-based assay. In addition, as revealed from the predicted binding model, further exploration of the binding interactions of this series of UKI-1 inhibitor bound at the S2 hydrophobic pocket would improve the potency and selectivity.

A number of hits identified from the VS which were not reported as serine protease inhibitors also showed inhibition in the TMPRSS2 assay with 50%-70% maximal efficacy. Although these compounds did not show viral entry inhibition in the PP assay, most likely due to their weak potency and efficacy, they represent a diversity of scaffolds that may serve as starting points for further structure-based optimization for potent and selective inhibitors against TMPRSS2. Most of these inhibitors possess a benzoamidinium head group, reiterating a key role of the functional head group for effective binding and inhibition. Indeed, a number of non-amidinium-based compounds which fit well in the active site of TMPRSS2 were selected from VS and tested in the enzyme assay, but none of them showed efficient activity, indicating a challenging problem to design novel inhibitors against this serine protease. NCGC00487181 with a sulfonamide group is an interesting inhibitor. It is a structural analog of Celecoxib, a COX-2 inhibitor as nonsteroidal anti-inflammatory drug, which was used for the treatment of COVID-19 and was reported to weakly inhibit the main protease of the virus (Mpro).^49^ NCGC00487181 showed a potency of 3.49 µM of IC_50_ and 70% of efficacy on TMPRSS2 in the enzyme assay. However, it had significant fluorescent interference in the assay and was inactive in the PP assay, therefore, the mechanism of this interesting series of compounds on SARS-COV-2 viral inhibition need further investigation through an orthogonal assay platform. In addition, we also identified several quinol-like compounds with approximately 40% maximal efficacy in the TMPRSS2 enzyme assay (**Figure S8**). Although these compounds are not structurally interesting as drug candidates and their antiviral activity on SARS-COV-2 needs further validation, they are predominately occupied at the catalytic site of TMPRSS2 by catching H-binding interactions with the triad residue, therefore, may provide a template for hybrid design of novel inhibitors against TMPRSS2.

It is worth mentioning that the inhibitors identified from this VS campaign are expected to be non-covalent protease inhibitors. This is in part due to the mechanism of docking-based approach which is not capable of identifying a covalent binder. On the other hand, we were inclined to deprioritize the traditional covalent protease inhibitors in the VS in order to identify novel and non-covalent chemotypes. Compared to the covalent inhibitors such as camostat and nafamostat with a reactive functional group to the highly conserved catalytic site,^50^ the non-covalent inhibitors are less chemically and metabolically reactive; therefore, these are more advantageous and attractive in the design of selective inhibitors. A number of non-covalent inhibitors of serine proteases thrombin and factor Xa have been reported with better selectivity.^51^ As revealed from the binding models of these identified inhibitors, TMPRSS2 showed different structural features and binding specificity at the distal hydrophobic pocket, which makes it a promising target for structure-based design and chemistry lead optimization to achieve selectivity and drug-like properties as drug candidates for the treatment of COVID-19.

## Supporting Information

Details of TMPRSS structural model and MD simulations; Pharmacophore model of TMPRSS2 used for virtual screening; Experimental data of identified TMPRSS2 inhibitors, their predicted binding models and calculated chemical properties.

## ACKNOWLEDGMENTS

This work was supported by the Intramural Research Programs of the National Center for Advancing Translational Sciences, National Institutes of Health.

## REFERENCES

1. Lai, C. C., Shih, T. P., Ko, W. C., Tang, H. J., Hsueh, P. R., Severe acute respiratory syndrome coronavirus 2 (SARS-CoV-2) and coronavirus disease-2019 (COVID-19): The epidemic and the challenges. Int J Antimicrob Agents 2020, 55 (3), 105924.

2. Lu, R., Zhao, X., Li, J., Niu, P., Yang, B., Wu, H., Wang, W., Song, H., Huang, B., Zhu, N., Bi, Y., Ma, X., Zhan, F., Wang, L., Hu, T., Zhou, H., Hu, Z., Zhou, W., Zhao, L., Chen, J., Meng, Y., Wang, J., Lin, Y., Yuan, J., Xie, Z., Ma, J., Liu, W. J., Wang, D., Xu, W., Holmes, E.C., Gao G. F., Wu, G., Chen, W., Shi, W., Tan, W., Genomic characterisation and epidemiology of 2019 novel coronavirus: implications for virus origins and receptor binding. Lancet 2020, 395 (10224), 565–574.

3. Zhou, P., Yang, X. L., Wang, X. G., Hu, B., Zhang, L., Zhang, W., Si, H. R., Zhu, Y., Li, B., Huang, C. L., Chen, H. D., Chen, J., Luo, Y., Guo, H., Jiang, R. D., Liu, M. Q., Chen, Y., Shen, X. R., Wang, X., Zheng, X. S., Zhao, K., Chen, Q. J., Deng, F., Liu, L. L., Yan, B., Zhan, F. X., Wang, Y. Y., Xiao, G. F., Shi, Z. L., A pneumonia outbreak associated with a new coronavirus of probable bat origin. Nature 2020, 579 (7798), 270–273.

4. Vinayagam, S., Sattu, K., SARS-CoV-2 and coagulation disorders in different organs. Life Sci 2020, 260, 118431.

5. Ortega-Paz, L., Capodanno, D., Montalescot, G., Angiolillo, D. J., COVID-19 Associated Thrombosis and Coagulopathy: Review of the Pathophysiology and Implications for Antithrombotic Management. J Am Heart Assoc 2020, e019650.

6. Lu, H., Drug treatment options for the 2019-new coronavirus (2019-nCoV). Biosci Trends 2020, 14 (1), 69–71.

7. Sheahan, T. P., Sims, A. C., Leist, S. R., Schafer, A., Won, J., Brown, A. J., Montgomery, S. A., Hogg, A., Babusis, D., Clarke, M. O., Spahn, J. E., Bauer, L., Sellers, S., Porter, D., Feng, J. Y., Cihlar, T., Jordan, R., Denison, M. R., Baric, R. S., Comparative therapeutic efficacy of remdesivir and combination lopinavir, ritonavir, and interferon beta against MERS-CoV. Nat Commun 2020, 11 (1), 222.

8. Pillaiyar, T., Meenakshisundaram, S., Manickam, M., Recent discovery and development of inhibitors targeting coronaviruses. Drug Discov Today 2020, 25 (4), 668–688.

9. Wang, M., Cao, R., Zhang, L., Yang, X., Liu, J., Xu, M., Shi, Z., Hu, Z., Zhong, W., Xiao, G., Remdesivir and chloroquine effectively inhibit the recently emerged novel coronavirus (2019-nCoV) in vitro. Cell Res 2020, 30 (3), 269–271.

10. Liu, J., Cao, R., Xu, M., Wang, X., Zhang, H., Hu, H., Li, Y., Hu, Z., Zhong, W., Wang, M., Hydroxychloroquine, a less toxic derivative of chloroquine, is effective in inhibiting SARS-CoV-2 infection in vitro. Cell Discov 2020, 6, 16.

11. Velthuis, A.J., van den Worm, S.H., Sims, A.C., Baric, R.S., Snijder, E. J., van Hemert, M.J., Zn(2+) inhibits coronavirus and arterivirus RNA polymerase activity in vitro and zinc ionophores block the replication of these viruses in cell culture. PLoS Pathog 2010, 6 (11), e1001176.

12. Derwand, R., Scholz, M., Does zinc supplementation enhance the clinical efficacy of chloroquine/hydroxychloroquine to win today’s battle against COVID-19? Med Hypotheses 2020, 142, 109815.

13. Touret, F., Gilles, M., Barral, K., Nougairede, A., van Helden, J., Decroly, E., de Lamballerie, X., Coutard, B., In vitro screening of a FDA approved chemical library reveals potential inhibitors of SARS-CoV-2 replication. Sci Rep 2020, 10 (1), 13093.

14. Choy, K. T., Wong, A. Y., Kaewpreedee, P., Sia, S. F., Chen, D., Hui, K. P. Y., Chu, D. K. W., Chan, M. C. W., Cheung, P. P., Huang, X., Peiris, M., Yen, H. L., Remdesivir, lopinavir, emetine, and homoharringtonine inhibit SARS-CoV-2 replication in vitro. Antiviral Res 2020, 178, 104786.

15. Eastman, R. T., Roth, J. S., Brimacombe, K. R., Simeonov, A., Shen, M., Patnaik, S., Hall, M. D., Remdesivir: A Review of Its Discovery and Development Leading to Emergency Use Authorization for Treatment of COVID-19. ACS Cent Sci 2020, 6 (5), 672–683.

16. Li, G., De Clercq, E., Therapeutic options for the 2019 novel coronavirus (2019-nCoV). Nat Rev Drug Discov 2020, 19 (3), 149–150.

17. Li, W., Moore, M. J., Vasilieva, N., Sui, J., Wong, S. K., Berne, M. A., Somasundaran, M., Sullivan, J. L., Luzuriaga, K., Greenough, T. C., Choe, H., Farzan, M., Angiotensin-converting enzyme 2 is a functional receptor for the SARS coronavirus. Nature 2003, 426 (6965), 450–4.

18. Glowacka, I., Bertram, S., Muller, M. A., Allen, P., Soilleux, E., Pfefferle, S., Steffen, I., Tsegaye, T. S., He, Y., Gnirss, K., Niemeyer, D., Schneider, H., Drosten, C., Pohlmann, S., Evidence that TMPRSS2 activates the severe acute respiratory syndrome coronavirus spike protein for membrane fusion and reduces viral control by the humoral immune response. J Virol 2011, 85 (9), 4122–34.

19. Shirato, K., Kawase, M., Matsuyama, S., Middle East respiratory syndrome coronavirus infection mediated by the transmembrane serine protease TMPRSS2. J Virol 2013, 87 (23), 12552–61.

20. Yamamoto, M., Matsuyama, S., Li, X., Takeda, M., Kawaguchi, Y., Inoue, J. I., Matsuda, Z., Identification of Nafamostat as a Potent Inhibitor of Middle East Respiratory Syndrome Coronavirus S Protein-Mediated Membrane Fusion Using the Split-Protein-Based Cell-Cell Fusion Assay. Antimicrob Agents Chemother 2016, 60 (11), 6532–6539.

21. Hoffmann, M., Kleine-Weber, H., Schroeder, S., Kruger, N., Herrler, T., Erichsen, S., Schiergens, T. S., Herrler, G., Wu, N. H., Nitsche, A., Muller, M. A., Drosten, C., Pohlmann, S., SARS-CoV-2 Cell Entry Depends on ACE2 and TMPRSS2 and Is Blocked by a Clinically Proven Protease Inhibitor. Cell 2020, 181 (2), 271–280 e8.

22. Bestle, D., Heindl, M. R., Limburg, H., Van Lam van, T., Pilgram, O., Moulton, H., Stein, D. A., Hardes, K., Eickmann, M., Dolnik, O., Rohde, C., Klenk, H. D., Garten, W., Steinmetzer, T., Bottcher-Friebertshauser, E., TMPRSS2 and furin are both essential for proteolytic activation of SARS-CoV-2 in human airway cells. Life Sci Alliance 2020, 3 (9).

23. Hoffmann, M., Hofmann-Winkler, H., Smith, J. C., Kruger, N., Sorensen, L. K., Sogaard, O. S., Hasselstrom, J. B., Winkler, M., Hempel, T., Raich, L., Olsson, S., Yamazoe, T., Yamatsuta, K., Mizuno, H., Ludwig, S., Noe, F., Sheltzer, J. M., Kjolby, M., Pohlmann, S., Camostat mesylate inhibits SARS-CoV-2 activation by TMPRSS2-related proteases and its metabolite GBPA exerts antiviral activity. bioRxiv 2020.

24. Naik, B., Gupta, N., Ojha, R., Singh, S., Prajapati, V. K., Prusty, D., High throughput virtual screening reveals SARS-CoV-2 multi-target binding natural compounds to lead instant therapy for COVID-19 treatment. Int J Biol Macromol 2020, 160, 1–17.

25. Shrimp, J. H., Kales, S. C., Sanderson, P. E., Simeonov, A., Shen, M., Hall, M. D., An Enzymatic TMPRSS2 Assay for Assessment of Clinical Candidates and Discovery of Inhibitors as Potential Treatment of COVID-19. bioRxiv 2020.

26. Gorshkov, K., Chen, C. Z., Bostwick, R., Rasmussen, L., Xu, M., Pradhan, M., Tran, B. N., Zhu, W., Shamim, K., Huang, W., Hu, X., Shen, M., Klumpp-Thomas, C., Itkin, Z., Shinn, P., Simeonov, A., Michael, S., Hall, M. D., Lo, D. C., Zheng, W., The SARS-CoV-2 cytopathic effect is blocked with autophagy modulators. bioRxiv 2020.

27. Roy, A., Kucukural, A., Zhang, Y., I-TASSER: a unified platform for automated protein structure and function prediction. Nat Protoc 2010, 5 (4), 725–38.

28. Herter, S., Piper, D. E., Aaron, W., Gabriele, T., Cutler, G., Cao, P., Bhatt, A. S., Choe, Y., Craik, C. S., Walker, N., Meininger, D., Hoey, T., Austin, R. J., Hepatocyte growth factor is a preferred in vitro substrate for human hepsin, a membrane-anchored serine protease implicated in prostate and ovarian cancers. Biochem J 2005, 390 (Pt 1), 125–36.

29. Guertin, K. R., Gardner, C. J., Klein, S. I., Zulli, A. L., Czekaj, M., Gong, Y., Spada, A. P., Cheney, D. L., Maignan, S., Guilloteau, J. P., Brown, K. D., Colussi, D. J., Chu, V., Heran, C. L., Morgan, S. R., Bentley, R. G., Dunwiddie, C. T., Leadley, R. J., Pauls, H. W., Optimization of the beta-aminoester class of factor Xa inhibitors. Part 2: Identification of FXV673 as a potent and selective inhibitor with excellent In vivo anticoagulant activity. Bioorg Med Chem Lett 2002, 12 (12), 1671–4.

30. Zeslawska, E., Schweinitz, A., Karcher, A., Sondermann, P., Sperl, S., Sturzebecher, J., Jacob, U., Crystals of the urokinase type plasminogen activator variant beta(c)-uPAin complex with small molecule inhibitors open the way towards structure-based drug design. J Mol Biol 2000, 301 (2), 465–75.

31. Iwasaka, S., Shono, Y., Tokuda, K., Nakashima, K., Yamamoto, Y., Maki, J., Nagasaki, Y., Shimono, N., Akahoshi, T., Taguchi, T., Clinical improvement in a patient with severe coronavirus disease 2019 after administration of hydroxychloroquine and continuous hemodiafiltlation with nafamostat mesylate. J Infect Chemother 2020, 26 (12), 1319–1323.

32. Zhou, Y., Vedantham, P., Lu, K., Agudelo, J., Carrion, R., Jr., Nunneley, J. W., Barnard, D., Pohlmann, S., McKerrow, J. H., Renslo, A. R., Simmons, G., Protease inhibitors targeting coronavirus and filovirus entry. Antiviral Res 2015, 116, 76–84.

33. Yuksel, M., Okajima, K., Uchiba, M., Okabe, H., Gabexate mesilate, a synthetic protease inhibitor, inhibits lipopolysaccharide-induced tumor necrosis factor-alpha production by inhibiting activation of both nuclear factor-kappaB and activator protein-1 in human monocytes. J Pharmacol Exp Ther 2003, 305 (1), 298–305.

34. Molecular Operating Environment (MOE) 2019.01. Chemical Computing Group ULC, Montreal, QC, Canada.

35. Trott, O., Olson, A. J., AutoDock Vina: improving the speed and accuracy of docking with a new scoring function, efficient optimization, and multithreading. J Comput Chem 2010, 31 (2), 455–61.

36. Case, D. A., Ben-Shalom, I. Y., Brozell, S.R., D.S., C., T.E., C. I., Cruzeiro, V.W.D, Darden, T. A., Duke, R. E., Ghoreishi, D., Simmerling CL; Wang J; Luo R; Merz KM; Wang B; Pearlman DA; Crowley M; Tsui V; Gohlke H; Mongan J; Hornak V; Cui G; Beroza P; Schafmeister C; Walker, R.C., Wei, H., Wolf, R. M., Wu, X., Xiao, L., York, D. M., Kollman, P. A., AMBER18. University of California, San Francisco: 2018.

37. Hu, X., Zhang, Y. Q., Lee, O. W., Liu, L., Tang, M., Lai, K., Boxer, M. B., Hall, M. D., Shen, M., Discovery of novel inhibitors of human galactokinase by virtual screening. J Comput Aided Mol Des 2019, 33 (4), 405–417.

38. Hawkins, P. C., Skillman, A. G., Nicholls, A., Comparison of shape-matching and docking as virtual screening tools. J Med Chem 2007, 50 (1), 74–82.

39. Chen, C. Z., Xu, M., Pradhan, M., Gorshkov, K., Petersen, J., Straus, M. R., Zhu, W., Shinn, P., Guo, H., Shen, M., Klumpp-Thomas, C., Michael, S. G., Zimmerberg, J., Zheng, W., Whittaker, G. R., Identifying SARS-CoV-2 entry inhibitors through drug repurposing screens of SARS-S and MERS-S pseudotyped particles. bioRxiv 2020.

40. Millet, J. K., Whittaker, G. R., Murine Leukemia Virus (MLV)-based Coronavirus Spike-pseudotyped Particle Production and Infection. Bio Protoc 2016, 6 (23).

41. Rensi, S., Altman, R. B., Liu, T., Lo, Y. C., McInnes, G., Derry, A., Keys, A., Homology Modeling of TMPRSS2 Yields Candidate Drugs That May Inhibit Entry of SARS-CoV-2 into Human Cells. ChemRxiv 2020.

42. Hedstrom, L., Serine protease mechanism and specificity. Chem Rev 2002, 102 (12), 4501–24.

43. Hempel, T., Raich, L., Olsson, S., Azouz, N., Klingler, A., Rothenberg, M., Noe, F., Molecular mechanism of SARS-CoV-2 cell entry inhibition via TMPRSS2 by Camostat and Nafamostat mesylate. bioRxiv 2020, doi: https://doi.org/10.1101/2020.07.21.214098.

44. Sun, Y. J., Velez, G., Parsons, D., Bassuk, A. G., Mahajan, V., TMPRSS2 structure-phylogeny repositions Avoralstat for SARS-1 CoV-2 prophylaxis in mice. bioRxiv 2021.

45. Zhu, W., Xu, M., Chen, C. Z., Guo, H., Shen, M., Hu, X., Shinn, P., Klumpp-Thomas, C., Michael, S. G., Zheng, W., Identification of SARS-CoV-2 3CL Protease Inhibitors by a Quantitative High-throughput Screening. bioRxiv 2020.

46. Guertin, K. R., Choi, Y. M., The discovery of the Factor Xa inhibitor otamixaban: from lead identification to clinical development. Curr Med Chem 2007, 14 (23), 2471–81.

47. Liao, Y., Bottcher, H., Harting, J., Greiner, H., van Amsterdam, C., Cremers, T., Sundell, S., Marz, J., Rautenberg, W., Wikstrom, H., New selective and potent 5-HT(1B/1D) antagonists: chemistry and pharmacological evaluation of N-piperazinylphenyl biphenylcarboxamides and biphenylsulfonamides. J Med Chem 2000, 43 (3), 517–25.

48. Sturzebecher, J., Vieweg, H., Steinmetzer, T., Schweinitz, A., Stubbs, M. T., Renatus, M., Wikstrom, P., 3-Amidinophenylalanine-based inhibitors of urokinase. Bioorg Med Chem Lett 1999, 9 (21), 3147–52.

49. Gimeno, A., Mestres-Truyol, J., Ojeda-Montes, M. J., Macip, G., Saldivar-Espinoza, B., Cereto-Massague, A., Pujadas, G., Garcia-Vallve, S., Prediction of Novel Inhibitors of the Main Protease (M-pro) of SARS-CoV-2 through Consensus Docking and Drug Reposition. Int J Mol Sci 2020, 21 (11).

50. Sun, W., Zhang, X., Cummings, M. D., Albarazanji, K., Wu, J., Wang, M., Alexander, R., Zhu, B., Zhang, Y., Leonard, J., Lanter, J., Lenhard, J., Targeting enteropeptidase with reversible covalent inhibitors to achieve metabolic benefits. J Pharmacol Exp Ther 2020.

51. Sanderson, P. E., Small, noncovalent serine protease inhibitors. Med Res Rev 1999, 19 (2), 179–97.

