## Supplementary material for "Discovery of TMPRSS2 inhibitors from virtual screening": Supplemetal Figures

**Discovery of TMPRSS2 inhibitors from virtual screening as potential treatment of COVID-19**

### Supplementary Information

### Table of Contents

- **Figure S-1** TMPRSS2 structural model superimposed with templates (pg. S-2)
- **Figure S-2** RMSF plot of TMPRSS2 in MD simulations (pg. S-3)
- **Figure S-3** H-bond plot of TMPRSS2 inhibitor binding in MD simulations (pg. S-4)
- **Figure S-4** Pharmacophore model used for TMPRSS2 virtual screening (pg. S-5)
- **Figure S-5** Activities of identified inhibitors in TMPRSS2 biochemical assay (pg. S-6)
- **Figure S-6** Activities of TMPRSS2 inhibitors in SARS-CoV-2 PP entry assay (pg. S-7)
- **Figure S-7** Predicted binding models of TMPRSS2 inhibitors (pg. S-8)
- **Figure S-8** Quinol-like hits identified from virtual screening (pg. S-9)
- **Table S-1** Chemical properties of TMPRSS2 inhibitors in this study (pg. S-10)

Figure S-1. TMPRSS2 structural model (blue) superimposed with template structure hepsin (magenta, PDB 1Z8G) and two related serine proteases Coagulation Factor Xa (brown, PDB 1KSN) and Urokinase (grey, PDB 1F92).

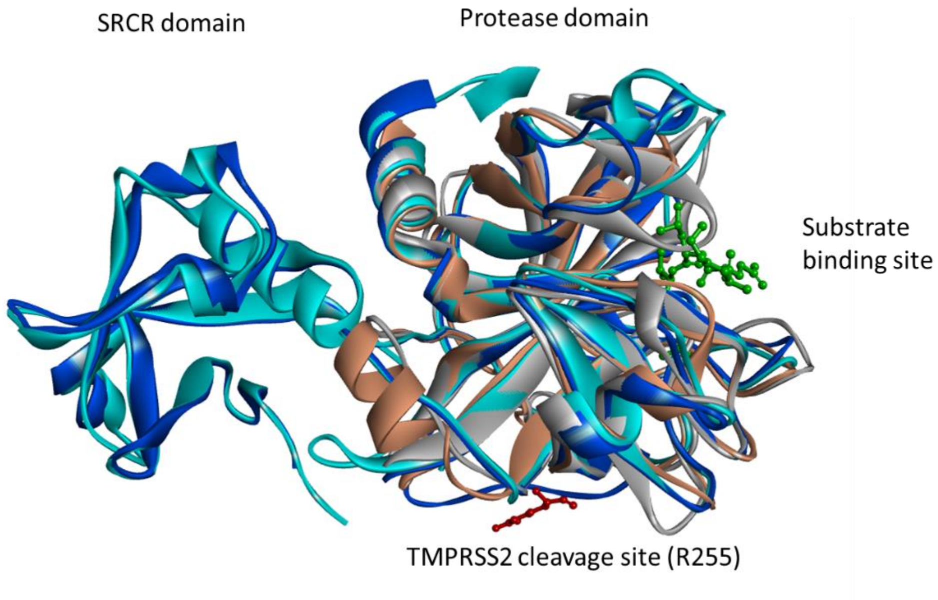

Figure S-2. RMSF plot of TMPRSS2 with the apo and inhibitor-bound complexes in the MD simulations.

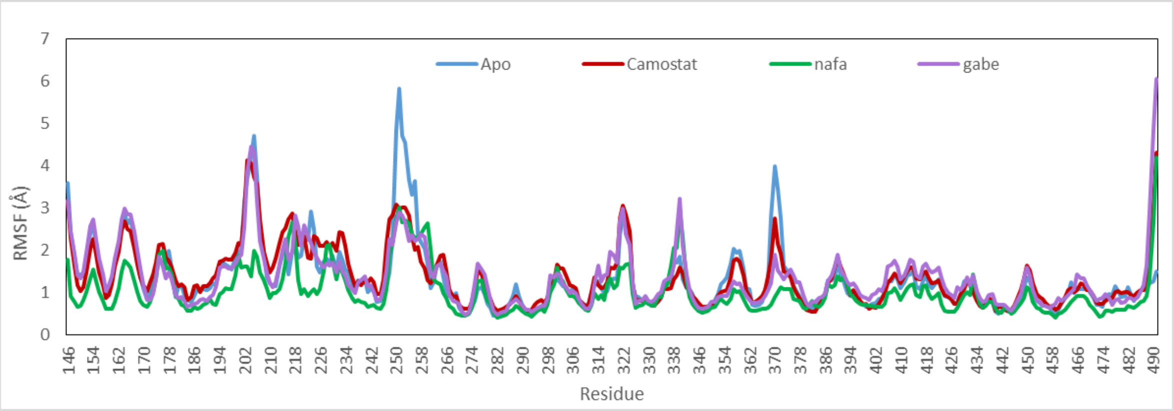

Figure S-3. Plot of H-bond interactions between the guanidinium head group of TMPRSS2 inhibitors bound and Asp239 in the 10-ns MD simulations.

**Camostat**

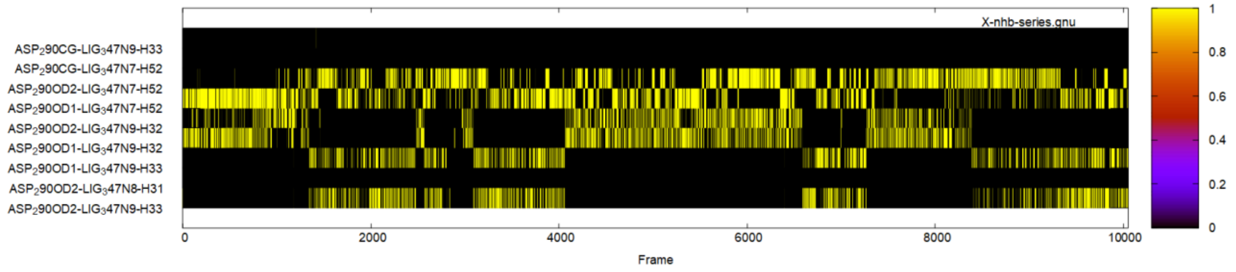


**Nafamostat**

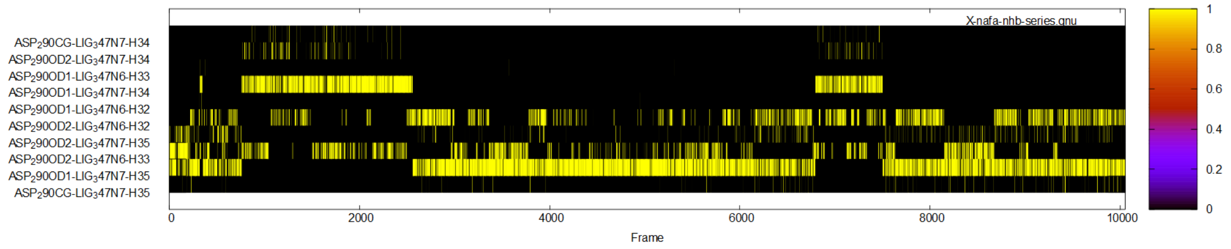


**Gabexate**

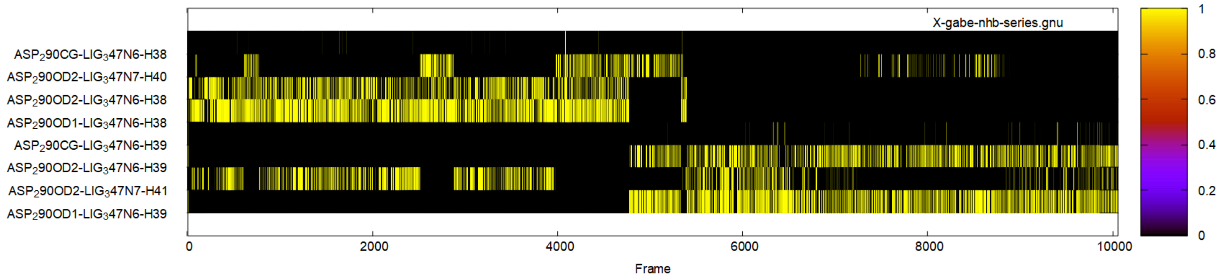

Figure S-4. Pharmacophore model used for TMPRSS2 virtual screening. The pharmacophore models were generated based on the predicted binding interactions of TMPRSS2 with inhibitor camostat (red), nafamostat (green) and gabexate (dark yellow) using MOE. Four pharmacophoric features were included: 1) an Don2 projected H-bond donor feature placed on the sidechain of Asp435 in the S1 pocket; 2) an Acc2 projected H-bond acceptor feature placed on the N atom of sidechain of Gln438; 3) an hydrophobic centroids Hyd feature matching hydrophobic interactions at the S1' hydrophobic region mainly formed by Val275, Val280, and Leu302; 4) an Don2 projected H-bond donor feature placed on the sidechain of Glu299.

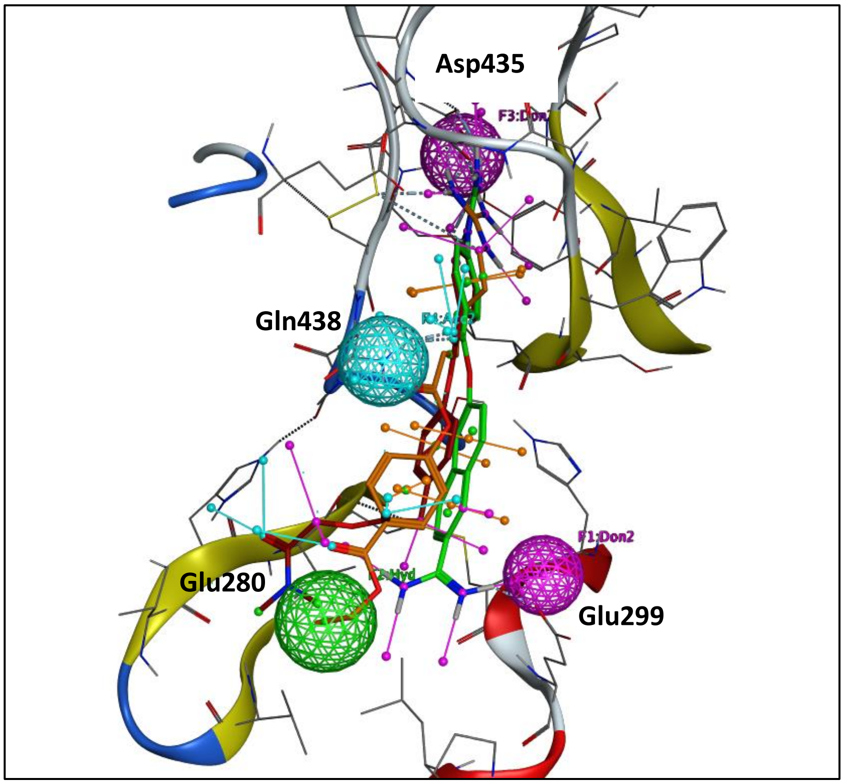

Figure S-5. Activities of identified inhibitors in the TMPRSS2 enzyme assay and counter screen.

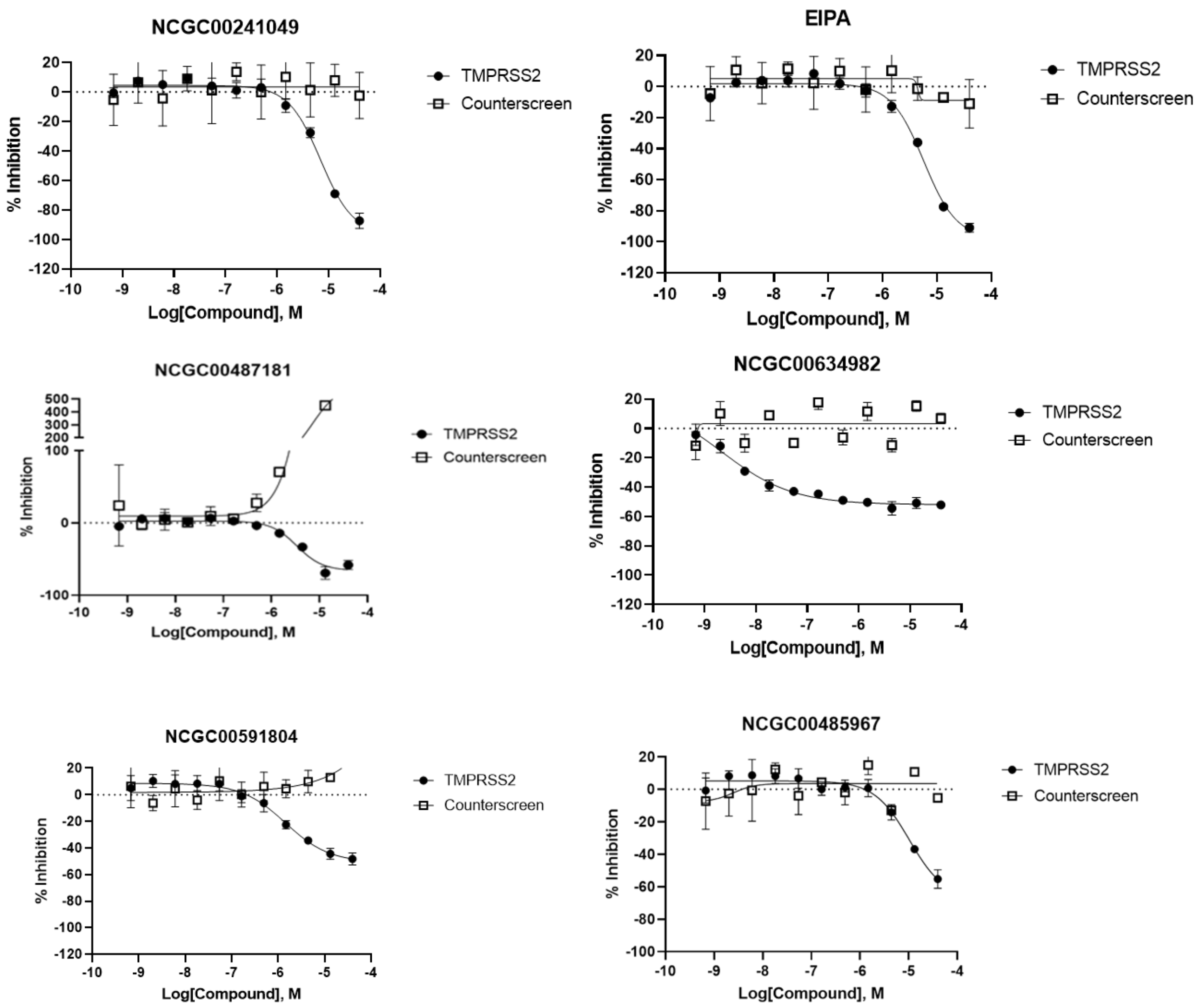

Figure S-6. Activities of TMPRSS2 inhibitors Avoralastat, PCI-27483 and Antipain in the SARS-COV-2 PP entry assay and counter screen.

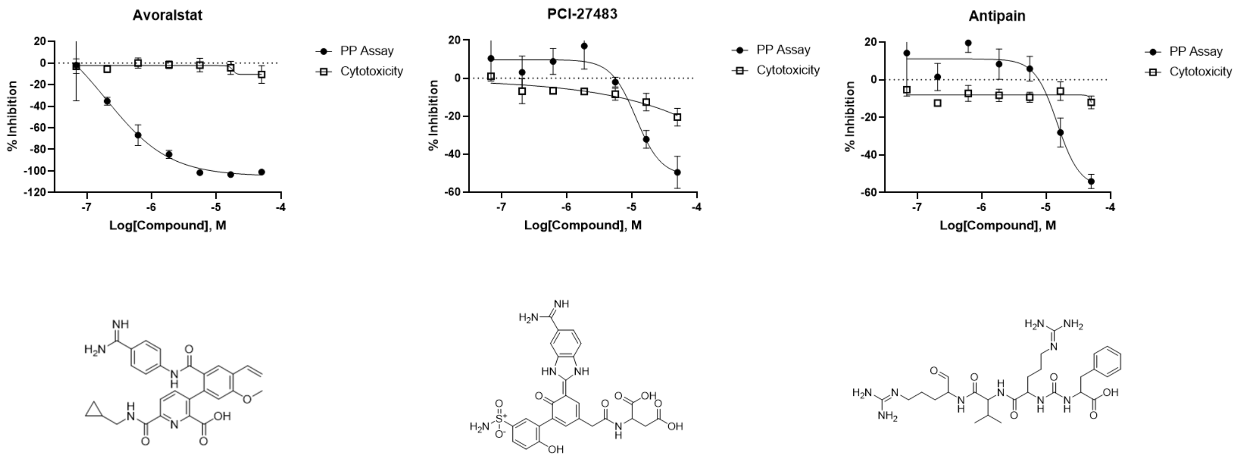

Figure S-7. Predicted binding models of TMPRSS2 inhibitors from VS. Protein surface is shown in hydrophobicity and small molecules are shown in sticks. Two binding models of otamixaban were predicted from docking analysis.

Otamixaban NCGC00591804 NCGC00487181

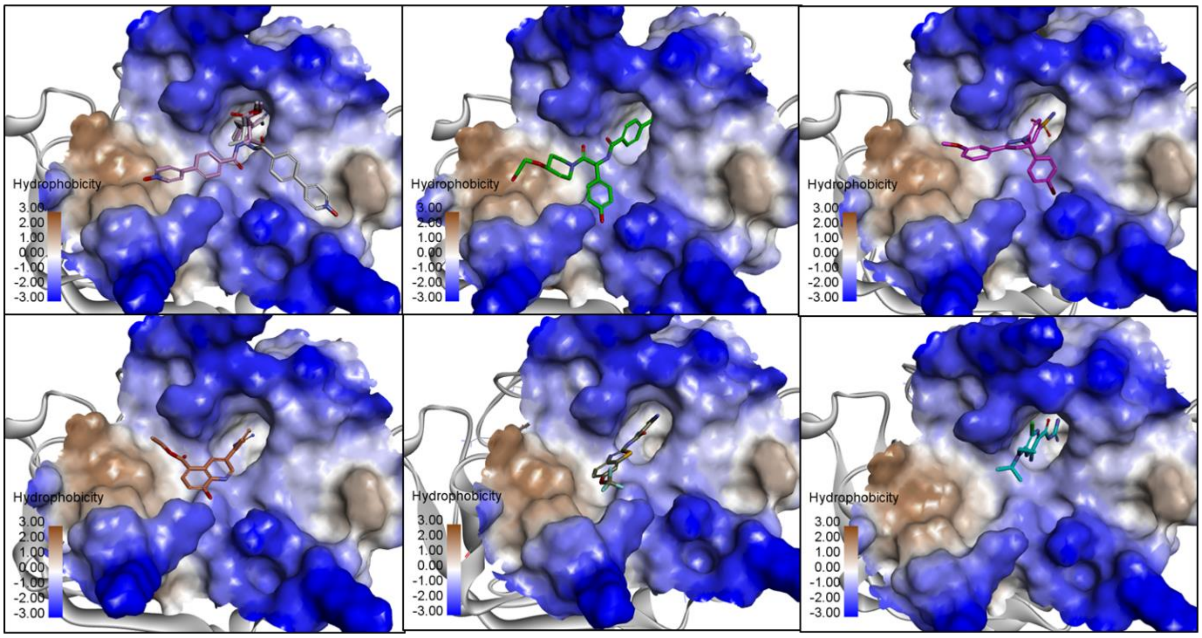


NCGC00241049 NCGC00634982 NCGC00387094

Figure S-8. Quinol-like hits identified from virtual screening and activities tested in the TMPRSS2 enzyme assay.

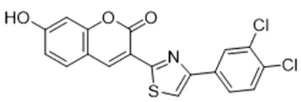

NCGC00138440
(1.96 uM, 46.7%)

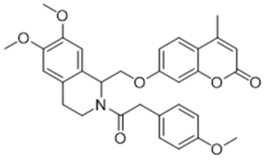

NCGC00104913
(40uM, 87.6%)

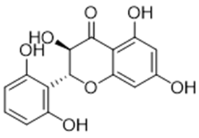

NCGC00385208
(27.7uM, 40.9%)

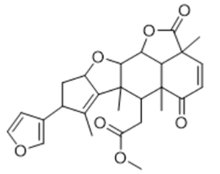

NCGC00390383
(35 uM, 32.7%)

**Table S-1.** Chemical properties of TMPRSS2 inhibitors in this study.

| NCGC ID | Name | MW | HB-Acc | HB-Don | LogP | TPSA |
|---|---|---|---|---|---|---|
| NCGC00160398 | Nafamostat Mesylate | 347.37 | 9 | 4 | 2.36 | 141.55 |
| NCGC00167526 | Camostat Mesylate | 398.41 | 5 | 5 | 1.37 | 136.55 |
| NCGC00025297 | Gabexate mesylate | 321.37 | 5 | 4 | 1.99 | 116.24 |
| NCGC00378763 | Otamixaban | 446.50 | 5 | 4 | 1.52 | 132.47 |
| NCGC00522442 | UKI-1 | 613.81 | 5 | 5 | 5.03 | 147.63 |
| NCGC00421880 | | 701.30 | 7 | 4 | 7.73 | 110.51 |
| NCGC00386945 | | 457.57 | 6 | 4 | 3.91 | 97.62 |
| NCGC00485967 | CBB1007 | 534.61 | 9 | 6 | -0.16 | 176.62 |
| NCGC00591804 | | 468.50 | 7 | 7 | -0.50 | 170.61 |
| NCGC00241049 | | 335.36 | 6 | 3 | 3.14 | 111.03 |
| NCGC00387094 | EIPA | 299.76 | 7 | 6 | 1.43 | 135.75 |
| NCGC00487181 | CID44216842 | 486.38 | 3 | 4 | 4.62 | 84.99 |
| NCGC00634982 | | 355.30 | 3 | 6 | 3.64 | 103.02 |
| NCGC00138440 | | 390.24 | 1 | 3 | 5.44 | 62.25 |
| NCGC00385208 | | 304.25 | 5 | 7 | 1.81 | 127.45 |
| NCGC00104913 | | 529.58 | 0 | 6 | 4.36 | 83.53 |
| NCGC00390383 | Nimbolide | 466.52 | 0 | 4 | 3.11 | 92.04 |
| NCGC00387860 | PCI-27483 | 596.57 | 12 | 12 | -2.02 | 282.49 |
| NCGC00522636 | Avoralstat | 513.54 | 7 | 6 | 2.40 | 172.06 |
| NCGC00390338 | Antipain | 604.70 | 15 | 10 | -2.95 | 288.55 |
